# Disturbance and landscape characteristics interactively drive dispersal strategies in continuous and fragmented metacommunities

**DOI:** 10.64898/2026.03.13.711669

**Authors:** Stav Gelber, Britta Tietjen, Felix May

**Affiliations:** Institute of Biology, Theoretical Ecology, Freie Universität Berlin, Berlin, Germany; Berlin-Brandenburg Institute of Advanced Biodiversity Research (BBIB), Berlin, Germany

**Keywords:** dispersal, fragmentation, disturbance, habitat amount, environmental autocorrelation, individual-based model

## Abstract

Habitat fragmentation, driven by human activities, disrupts habitat connectivity and alters ecological processes through geometric and demographic fragmentation effects. Dispersal plays a fundamental role in shaping the distribution, abundance, and persistence of species in modified landscapes. While previous research looked at the evolution of dispersal strategies at the species level, community-level dynamics remain underexplored. Species exhibit diverse dispersal strategies to persist in modified landscapes, yet predicting how these strategies interact at the community level requires a more integrated approach. This study employed an individual-based simulation model to explore how fragmentation and other landscape characteristics influence community-level dispersal strategies. We tested the effects of varying fragmentation levels, environmental autocorrelation, habitat amount, and disturbance levels on the emerging distribution of dispersal distances within a community in modified and continuous landscapes. We hypothesised that fragmentation and other spatial patterns would significantly shape community composition, favouring particular dispersal strategies under specific environmental conditions. The findings reveal that higher disturbance levels and greater habitat amount increased the community-weighted mean of dispersal distance, while fragmentation showed only minor variation. Additionally, low autocorrelation was associated with the highest community-weighted mean of dispersal distance. These results highlight the importance of considering community-level dynamics when predicting ecosystem responses to landscape modification. By clarifying how landscape structure and disturbance shape community-level dispersal strategies, this study advances our understanding of the mechanisms underlying species persistence and community structure in modified landscapes.

## Introduction

Major drivers of biodiversity change include climate change, invasive species, pollution, and land-use change. Anthropogenic land-use change, involving changes to the structure, distribution, or composition of natural environments, can negatively influence patterns of biodiversity and species abundance (Fischer & Lindenmayer, 2007; Foley et al., 2005; Sala et al., 2000). In addition to causing habitat loss and degradation, landscape modification can affect biodiversity most notably through habitat fragmentation. Habitat fragmentation is commonly a consequence of anthropogenic activities such as deforestation and urbanisation. However, the term ‘fragmentation’ is often used to mean different things. Some refer to habitat fragmentation (per se) as a specific difference in spatial configuration among landscapes with a given, constant habitat amount. Increasing fragmentation (per se) means that the constant habitat amount is provided by an increasing number of habitat fragments with decreasing average fragment sizes (Fahrig, 2003). Others consider fragmentation not only to be the subdivision of a constant habitat amount into smaller patches, but also as a more general dynamic process of landscape change that includes both habitat loss as well as changes in spatial configuration (Haddad et al., 2015). These differences in the definition of the term ‘fragmentation’ have been a source of confusion in the debate on whether fragmentation affects biodiversity positively or negatively (Valente et al., 2023). Importantly, in the context of this study, we refer to the former, more specific definition as ‘fragmentation’ while the latter, more general process including habitat loss as well as changes in configuration, is referred to as ‘landscape modification’.

Although the ecological consequences of habitat fragmentation are debated, particularly regarding whether its effects are positive or negative for biodiversity (Fahrig et al., 2019; Fletcher et al., 2018; Valente et al., 2023), there is broad agreement that both habitat loss and fragmentation can have significant ecological impacts on species composition and ecosystem functioning (Haddad et al., 2015). By changing the spatial configuration of habitats, fragmentation affects the mean size of habitat fragments, levels of isolation, and the amount of edge area. To better understand the ecological consequences of habitat loss and fragmentation, it is essential to explore the processes that determine how populations and communities respond to altered spatial patterns. These include demographic processes, species interactions, genetic processes, and, critically, dispersal and connectivity.

Dispersal is a fundamental ecological process that shapes the spatial distribution and abundance of species in their environment. Understanding dispersal strategies is essential for predicting species persistence in modified landscapes, particularly in metacommunity contexts where species distribution and diversity patterns at least partly depend on dispersal-driven dynamics (Leibold et al., 2004). A better understanding of the factors that drive the selection of dispersal distances within metacommunities is thus crucial for predicting how biodiversity will respond to landscape modification.

Accordingly, a key question in ecological research on dispersal is how habitat loss and fragmentation affect the performance of species and populations with different dispersal strategies and how this shapes trait distributions and (functional) diversity patterns of ecological communities. One prominent view, based on classical metapopulation theory, suggests that habitat loss and fragmentation should favour increased dispersal, since longer dispersal distances increase the (re)-colonisation rate of unoccupied habitat fragments and counteract local extinctions through the spatial rescue effect (Gotelli, 1991; Hanski, 1999). Some empirical studies examining short-term ecological responses support this prediction, showing that species with greater dispersal ability are favoured in modified landscapes, as observed in butterfly populations (Thomas, 2000) and ground beetle populations (de Vries et al., 1996). This is presumably due to the spatial rescue effect and the exploitation of ephemeral resources. Other research suggests the opposite, with landscape modification selecting for shorter dispersal distances. This opposing view is often linked to Darwin’s wind dispersal hypothesis, which posits that the costs of migration increase when a suitable habitat is rare. In particular, lower habitat amount and higher fragmentation reduce the chances of a propagule or offspring reaching a suitable habitat patch, especially with passive dispersal (Finand, Monnin, et al., 2024; Riba et al., 2009; Travis & Dytham, 1999). Under these conditions, long-distance dispersers risk arriving in unsuitable habitats with high chances of failing to establish. Accordingly, selection should favour shorter dispersal distances. Crucially, interactions between different factors, such as habitat amount and fragmentation, can jointly affect dispersal costs and strategies by altering the distance to suitable habitats (Travis & Dytham, 1999).

Empirical support for landscape modification selecting for shorter dispersal distances due to increased costs comes primarily from single-species evolutionary studies in plant seed dispersal (Riba et al., 2009) and ant populations (Finand, Monnin, et al., 2024). A single-species evolutionary simulation study also found that increasing fragmentation primarily selected for more dispersal to the natal site of sessile organisms, which will also reduce mean dispersal distances, while habitat amount had surprisingly weak effects in this study (Hovestadt et al., 2001). These conflicting findings highlight the context-dependency of dispersal strategies in modified landscapes, underscoring the need to consider additional ecological drivers that influence the distribution of suitable habitat and increase the selection pressure on dispersal distances.

In addition to landscape modification, the disturbance regime represents another major driver of metacommunity dynamics. Disturbance – typically defined as a discrete event that temporarily disrupts ecosystem structure, alters resource availability, or modifies the physical environment (Pickett & White, 1985) – can substantially reshape community structure and diversity. Events such as fire, storms, and deforestation can alter species distributions and drive the selection of dispersal strategies. Theoretical and empirical studies indicate that disturbance typically favours longer dispersal distances, as species with greater dispersal ability are more likely to recolonise disturbed areas (Alvarez-Buylla & Martínez-Ramos, 1990; Büchi & Vuilleumier, 2012; Treep et al., 2021; Willson, 1993).

One factor which very likely exerts selection pressure on dispersal distances is the spatial autocorrelation of environmental variables. In this context, high environmental autocorrelation refers to similar environmental conditions in adjacent and nearby sites, while low environmental autocorrelation refers to unpredictable changes in environmental variables across space (see Fig. 1). Accordingly, there is the clear expectation that high environmental autocorrelation should select for short dispersal distances, because propagules/juveniles that disperse over a short distance have a higher chance of finding similar environmental conditions to the sites in which their mother successfully reproduced. In contrast, low environmental autocorrelation is expected to exert low selection pressure on dispersal distances since chances to find suitable habitat are not or weakly related to distance from the natal site. However, there are surprisingly few studies on the selection of dispersal in landscapes with different levels of environmental autocorrelation. In a theoretical study of metacommunities in continuous landscapes, Büchi & Vuilleumier (2012) investigated the interacting effects of disturbance and environmental autocorrelation on dispersal distances. In contrast to the expectation formulated above, they found that high autocorrelation selects for longer dispersal distances, while low autocorrelation selects for short dispersal distances. This apparent contradiction can be easily explained by a specific technical detail of their simulation model. In their model, propagules have a chance to establish in the same site as their mother. This means that, in the scenarios with low autocorrelation, the mother’s site is the only site with a predictably higher chance for suitable environmental conditions. Or in other words, the low autocorrelation scenarios of Büchi and Vuilleumier (2012) correspond to a scenario with perfect autocorrelation at a distance of zero (the natal site), but low autocorrelation elsewhere. Of course, this exerts selection pressure to stay in the mother’s site.

**Figure 1.**
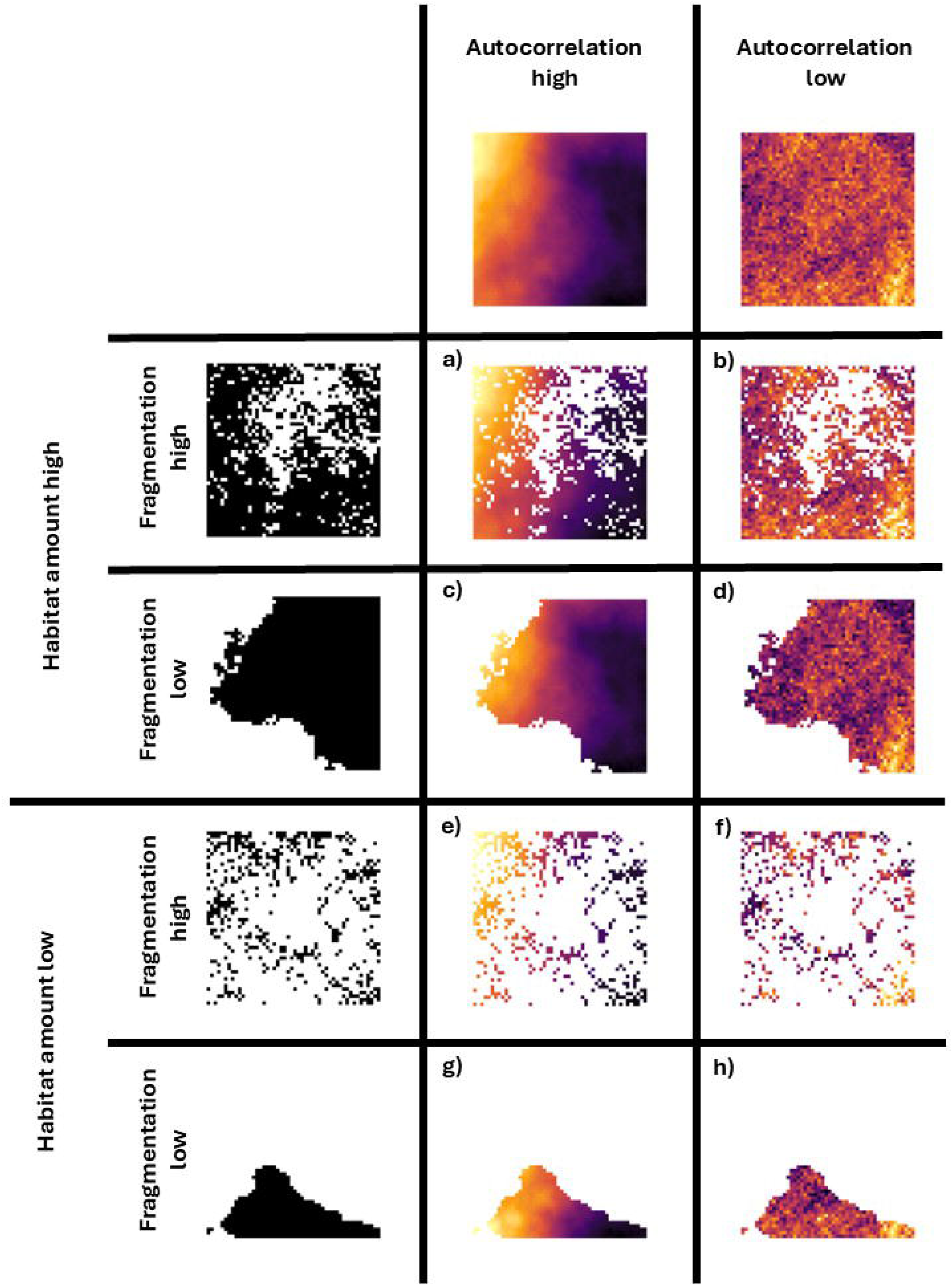
Illustration of the landscape configuration process. White cells represent matrix, colored cells represent habitat, and colour shade represents environmental values (0-1). Simulated landscapes (a–h) are generated from combinations of three properties: environmental autocorrelation (high in a, c, e, g; low in b, d, f, h), fragmentation level (high in a, b, e, f; low in c, d, g, h), and habitat amount (high in a–d; low in e–h). Each column represents a level of spatial environmental autocorrelation, while rows correspond to the combinations of fragmentation and habitat amount.

It should be noted that several studies cited here use the term “autocorrelation” in a way that differs from environmental autocorrelation. Specifically, in other studies, “autocorrelation” refers to the probability that a site with suitable habitat is next to another site with suitable habitat (e.g. Finand, Monnin, et al., 2024; Hovestadt et al., 2001; Travis & Dytham, 1999). In this way, this use of “autocorrelation” is effectively synonymous with the definition of fragmentation used here, since both refer to the spatial configuration of suitable habitat vs. unsuitable matrix, where low autocorrelation refers to high fragmentation (many small habitat fragments) and high autocorrelation refers to low fragmentation (few large habitat fragments). In the following, autocorrelation will always refer to environmental autocorrelation as explained above.

While previous studies have primarily focused on the response of dispersal at the population or metapopulation level, i.e. for single species (e.g., Clobert et al., 2004; Hanski, 2001), relatively less attention has been given to how different dispersal strategies are selected for at the community level, where multiple species interact and compete under different environmental conditions. We argue that a community-level approach is necessary to understand broader biodiversity impacts and patterns. Most evolutionary models allow for within-species variation in dispersal traits, where dispersal distances evolve plastically in response to environmental pressures (Travis et al., 2012). While dispersal traits can exhibit plasticity within species in response to environmental pressures, species tend to have characteristic dispersal strategies that are shaped by their evolutionary history and ecological constraints (Clobert et al., 2009). Importantly, a key difference between the evolution of populations and the selection from communities in response to fragmentation is at the temporal scale. Ecological dynamics at the community level, such as species extinctions and (re-)colonisations, occur much faster than evolutionary changes within populations. We therefore expect that landscape modification will impact community composition and diversity faster than populations can adapt through the evolution of altered dispersal strategies. Although plastic changes in dispersal can occur rapidly, they are often transient, maladaptive, or constrained (Valladares et al., 2007), and may not be comparable with the speed of community-level shifts in dispersal strategies.

Most previous studies investigated how single factors influence dispersal strategies. However, the interacting effects of environmental autocorrelation, disturbance, fragmentation, and habitat amount on the selection of dispersal distances remain underexplored, particularly at the community level. We aim to disentangle both the individual and interactive effects of these landscape characteristics on community dispersal strategies.

To explore these underlying mechanisms, we ran simulations of metacommunity dynamics in modified landscapes using an individual-based model. We simulate landscapes with varying levels of environmental autocorrelation, disturbance, fragmentation, and habitat amount. Specifically, we test the hypothesis that these factors alter the cost-benefit balance of dispersal, leading to shifts in the community weighted mean of dispersal distances. By explicitly considering community-level dispersal dynamics under different landscape and disturbance scenarios, our study provides new insights into the ecological mechanisms shaping dispersal strategies and biodiversity in fragmented landscapes.

## Methods

To explore the effects of the environmental variables on species dispersal strategies, we used a spatially explicit, individual-based model that simulates community dynamics in continuous and modified landscapes (Gelber et al., 2025). The model builds upon previous metacommunity frameworks (Gravel et al., 2006; Mouquet & Loreau, 2003; Rybicki & Hanski, 2013) and incorporates the processes of reproduction, mortality, competition, and dispersal. The simulated landscapes vary systematically in levels of environmental autocorrelation, habitat amount and fragmentation (Fig. 1).

The model includes two main entities: individuals and grid cells. Individuals are characterised by their species ID and location (XY-coordinates) and by their species-specific environmental optimum and niche breadth. Grid cells are characterised by their location, type (habitat or matrix), and, for habitat grid cells, their environmental value (between 0 and 1). That is, in contrast to other studies, we disentangle the general suitability of a grid cell to be a habitat and its environmental value. For simplicity, we make two broad assumptions: (i) Individuals are not able to move (i.e. we simulate sessile organisms such as plants), and (ii) individuals are not able to survive in the matrix cells. The spatial extent of the simulation is 200 x 200 cells, and each grid cell can host a single individual. Time is simulated with discrete steps (1000 time steps per simulation). In each time step, three processes occur sequentially: reproduction, death, and disturbance (optional). We use two types of landscapes in our simulations – modified and continuous. The simulation of modified landscapes is temporarily divided into two phases: initialisation and modified landscape (see the section “Continuous vs. Modified Landscape” below). Between these phases, landscape modification takes place as an instantaneous event using the “cookie-cutter” approach (May et al., 2019), and all individuals that are located in matrix cells after landscape modification instantly die. When simulating continuous landscapes, the landscape remains unmodified with 100% suitable habitat for the whole simulation (1000 time steps). Species richness and distribution patterns emerge from environmental heterogeneity of habitats, survival and recruitment probabilities, dispersal rates, and interspecific interactions via competition for space. The simulation model code, analysis scripts, and raw simulation output are archived on Zenodo (Gelber et al., 2026) and available on GitHub (https://github.com/Stavooo/Gelber_etal_2026_dispersal).

### Model Initialisation

We initialised the model with 1,000 possible species and generated 10,000 individuals, each randomly assigned to a species with equal probability, resulting in 10 individuals per species on average. Each species has a unique value for its maximum dispersal distance, environmental optimum, and ID. The birth and death rates, as well as niche breadth, are constant for all species (Table 1). The species-specific maximum dispersal distance values are randomly generated from a uniform distribution between 1 and 20. A single grid cell has a side length of 1. The environmental optimum values are generated as a sequence of 1,000 numbers (number of species) uniformly distributed between 0 and 1. Individuals are distributed randomly in habitat cells with environmental values that fall in the range of the species’ environmental optimum value (plus or minus its niche breadth).

**Table 1.** Model parameters used in experiments 1 to 3.

| Parameter name | Functionality | Accepted values | Exp. 1 | Exp. 2 | Exp. 3 |
| --- | --- | --- | --- | --- | --- |
| Grid side length | Determine simulation grid size | $\geq 1$ | 200 | 200 | 200 |
| Habitat amount | Proportion of habitat amount after modification | 0 - 1 | 0.1-0.9 | 0.1-0.9 | 0.15 |
| Spatial autocorrelation | Controls for landscape smoothness/roughness | 0 - 1 | 0.1-0.9 | 0.1-0.9 | 0.1 |
| Fragmentation | Level of fragmentation after modification | 0 - 1 | 0.1-0.9 | 0.1-0.9 | 0.7 |
| Population size | The initial number of individuals | $\geq 1$ | 10,000 | 10,000 | 10,000 |
| Species | Number of species in the species pool | $\geq 1$ | 1,000 | 1,000 | 1,000 |
| Niche breadth | Determine the range for accepted niche values | 0 - 1 | 0.2 | 0.2 | 0.2 |
| Reproduction rate | Proportion of successful reproduction events in each time step | 0 - 1 | 1 | 1 | 1 |
| Death rate | Proportion of death events in each time step | 0 - 1 | 0.1 | 0.1 | 0.1 |
| Dispersal distance | Mean distance for the dispersal kernel | $\geq 1$ | 1-20 | 1-20 | 1-20 |
| Carrying capacity | The number of individuals a cell can host | $\geq 1$ | 1 | 1 | 1 |
| Disturbance intensity | The rate at which the disturbance spreads | 0 - 1 | 0-0.4 | 0-0.4 | 0-0.4 |

We generated continuous and heterogeneous landscapes with a given environmental autocorrelation level (between 0 and 1, see below for details). The patterns of environmental values and landscape modifications are generated stochastically and thus differ between simulation runs. The simulation space is torus-shaped (i.e. we used periodic boundary conditions in the dispersal process) and therefore propagules closer to the grid edge have equal chances to reproduce successfully.

### Continuous vs. Modified Landscape

To simulate environmental heterogeneity across the landscape of 200 x 200 grid cells, we assigned an environmental value from the interval 0-1 to each habitat cell with varying levels of autocorrelation.

The environmental values are simulated by generating two-dimensional neutral landscapes using fractional Brownian motion (fBM) (Schlather et al., 2015; Sciaini et al., 2018; Travis & Dytham, 2004). When creating the landscapes, we controlled the spatial environmental autocorrelation using the Hurst coefficient H, which varies the ‘smoothness’ of the landscape (Hurst, 1951; Mandelbrot & Van Ness, 1968). The H values range from 0 (rugged landscape, with low environmental autocorrelation – Fig. 1 b, d, f, h) to 1 (smooth landscape, with high environmental autocorrelation – Fig. 1 a, c, e, g). The autocorrelation values we used in our experiments are shown in Table 1.

We conducted simulations in two types of landscapes: continuous and modified. In the continuous landscape, all grid cells remain habitat cells throughout the entire simulation (1000 time steps). For modified landscapes, the simulation begins with an initialisation phase in which the landscape remains continuous for the first 500 time steps. After this phase, the landscape is modified by changing a given proportion of the grid cells from habitat to matrix using a binarised landscape. The binarisation threshold (0–1) determines the proportion of habitat (habitat amount) in the final modified landscape. The Hurst coefficient governs the degree of fragmentation in this binary landscape – a low H value produces a highly fragmented landscape (Fig. 1 a, b, e, f), whereas a high H value results in a less fragmented landscape (Fig. 1 c, d, g, h) (Gelber et al., 2025; May et al., 2019). In our results, however, we invert the fragmentation values (Fragmentation level = 1 - H factor) for a more intuitive interpretation where low values represent a less fragmented landscape and high values represent a high level of fragmentation. The environmental values in the cells that remain as habitat are not changed during landscape modification (Fig. 1).

### Model Processes

#### Reproduction, dispersal, establishment, and death

During the reproduction process, the model loops through all individuals sequentially. In every time step, each individual produces one propagule. To determine the propagule’s target cell, we used a uniform dispersal kernel in a square around the focal cell with a unique maximum dispersal distance in the x and y directions for each species. If the target cell is a habitat cell that is not already occupied, the establishment probability of the propagule in the target cell is calculated following Gravel et al. (2006):

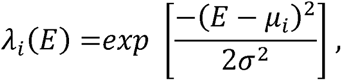

Where λi is the establishment probability of species i, E is the cell environmental value, μi is the species-specific environmental optimum for species i, and σ is the niche width. Accordingly, the establishment probability equals 1 if the propagule encounters its species-specific environmental optimum and decays exponentially (with a rate determined by the parameter σ) for values lower or higher than the optimum. Conversely, if the target cell is a matrix cell or already occupied, the propagule fails to establish and is considered dead. Additionally, the model includes the death process, where at each time step, individuals die at random with a fixed probability (0.1).

#### Disturbance

In experiments that include disturbance, we initiated a disturbance event that loosely mimicked a fire or a pest outbreak. At each time step, a disturbance event occurs with a set probability (0.25 disturbance rate), i.e. we assume the disturbance frequency to be fixed. During such an event, 3% of habitat cells are randomly selected as the initial disturbance points. The disturbance then spreads based on a predefined “disturbance spread rate”, affecting neighbouring cells in the four cardinal directions with a probability equal to this rate. This spread process continues iteratively until no additional cells are disturbed in a given iteration. Once all disturbed cells have been determined, all individuals within them are killed. That is, in our scenarios, we evaluate the effects of disturbance intensity (determined by the spread rate), but not of disturbance frequency. The environmental values of the grid cells are not modified by disturbances.

### Simulation experiments and model analysis

We investigated the effects of environmental autocorrelation and disturbance on the distribution of dispersal distances at the community level in continuous landscapes, and the effects of autocorrelation, disturbance, fragmentation level, and habitat amount in modified landscapes. Community-level dispersal distances were measured as the community weighted mean of dispersal distances (CWMDD) across all species, which is calculated as the average species dispersal distance weighted by the number of individuals of each species at a given time during the simulation. Additionally, we looked at two other response variables – species richness and the standard deviation of dispersal distances (SDDD) across species. We conducted three simulation experiments:

1. In the first experiment, we ran four separate simulation sets to isolate the individual effects of our key variables on CWMDD, species richness, and SDDD throughout the whole simulation (time series). Two simulations were conducted in a continuous landscape, where we independently varied environmental autocorrelation (five levels) and disturbance (five levels). The other two simulations were conducted in a modified landscape, where we independently varied fragmentation (five levels) and habitat amount (five levels) without disturbance and with autocorrelation set to 0.5.
2. In the second experiment, we employed a full factorial design and varied two variables simultaneously to explore how they interactively influence CWMDD, species richness, and SDDD at the end of simulations. We used three levels of autocorrelation and of disturbance in a continuous landscape, and additionally, in a separate simulation, three levels of habitat amount with two levels of fragmentation in a modified landscape.
3. In the third experiment, we employed a full factorial design to explore how the interaction between three variables affected our response variables at the end of simulations. We used three levels of autocorrelation, disturbance, and habitat amount in a modified landscape. We excluded the fragmentation level variable after identifying in previous experiments that its effect was comparably weak.

In all simulations, species differed only in their environmental optimum and maximum dispersal distances, while other species-level parameters, such as niche width, reproduction, and mortality rates, were kept equal. A full list of all model parameters used in these experiments can be found in Table 1. All simulation results are based on twenty repetitions for each parameter combination. Analyses were performed using R Statistical Software (R Core Team, 2023), and simulations were executed on the FU Berlin High-Performance computer, Curta (Bennett et al., 2020).

## Results

### Single variable effects

We assessed how independently modifying single environmental variables, while keeping the others constant, influences three community-level metrics: community-level mean dispersal distance (CWMDD), species richness, and the standard deviation of dispersal distances among species (SDDD) (Fig. 2). Accordingly, we consecutively varied environmental autocorrelation, disturbance spread rate, habitat fragmentation level, and habitat amount.

**Figure 2.**
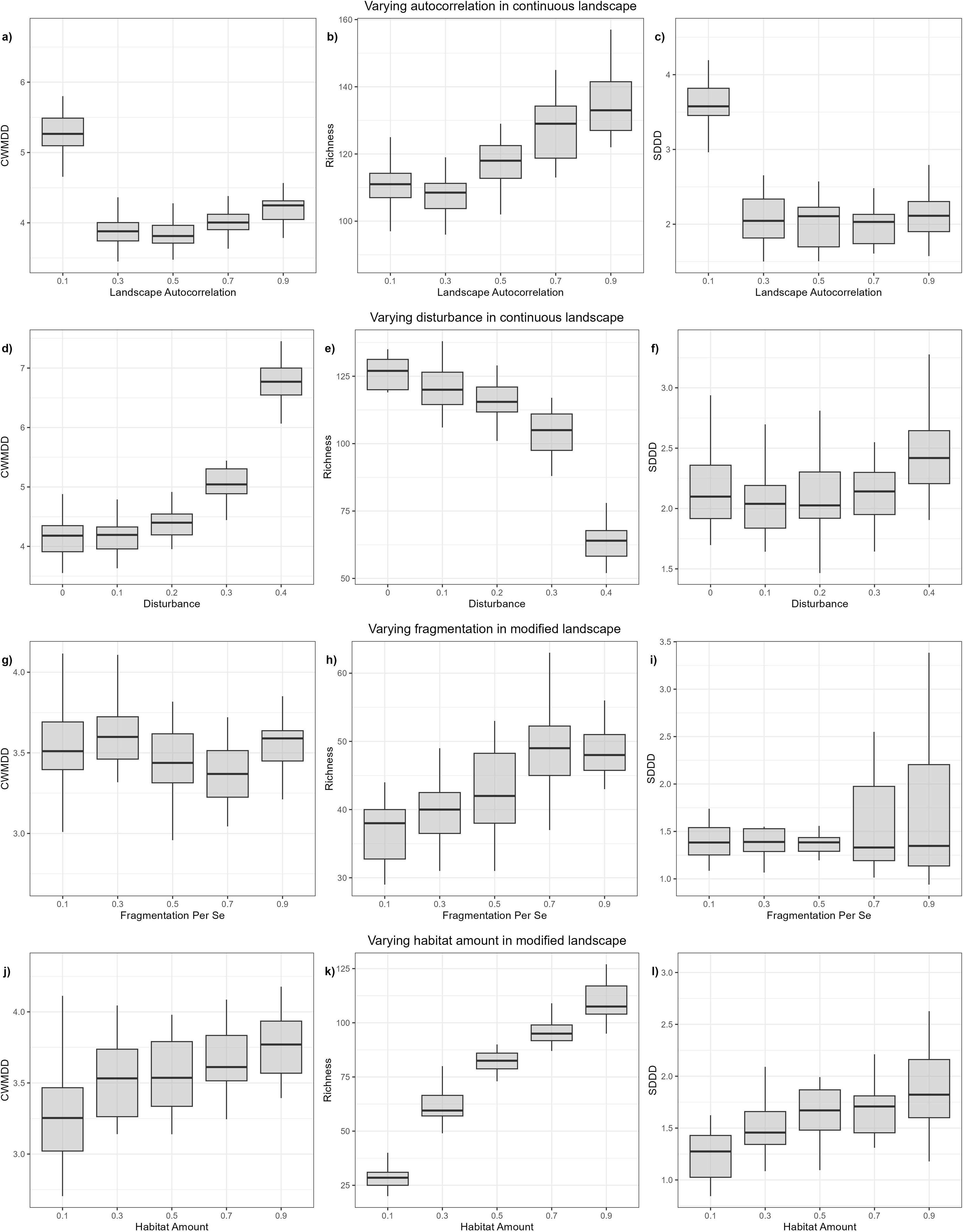
Simulation results showing the effects of independently modifying four landscape variables on three community-level response variables: Community-Weighted Mean Dispersal Distance (CWMDD, left column), species richness (middle column), and the standard deviation of dispersal distances (SDDD, right column). Rows represent different simulation scenarios, from top to bottom: varying autocorrelation, disturbance, fragmentation level, and habitat amount. Each parameter setting in every scenario was replicated 20 times for a total of 400 simulations, and all simulations ran for 1,000 time steps. Results were recorded at the end of each simulation. In the first two scenarios (autocorrelation and disturbance), the landscape remained continuous throughout the simulation. In the last two scenarios (fragmentation and habitat amount), the landscape was modified at time step 500 to introduce fragmentation. Disturbance occurred only in Scenario 2; all other scenarios had no disturbance. For Scenario 2, autocorrelation was held constant at 0.9. In Scenarios 3 and 4, autocorrelation was fixed at 0.5. In Scenario 3, the habitat amount was fixed at 0.2, and in Scenario 4, the fragmentation level was held constant at 0.7.

In the continuous landscape, CWMDD was highest in landscapes with low autocorrelation, declined with increasing autocorrelation, and reached its lowest values at intermediate levels. Still, the values at high environmental autocorrelation are only slightly higher than the values at intermediate autocorrelation (Fig. 2a). Species richness increased with higher levels of autocorrelation (Fig. 2b). SDDD was highest under low autocorrelation, whereas values at all other autocorrelation levels were comparatively low and similar (Fig. 2c).

With increasing disturbance spread rate, CWMDD increased while species richness decreased, indicating a negative relationship, as expected (Fig. 2d,e). SDDD was slightly higher under high disturbance but otherwise showed little variation across disturbance levels (Fig. 2f).

In the modified landscape, fragmentation levels caused only minor variation in CWMDD, with slightly higher values at intermediate fragmentation. However, these differences were small, and no consistent pattern emerged (Fig. 2g). Species richness increased with increasing fragmentation (Fig. 2h), while the median values of SDDD remained largely similar across fragmentation levels, the variation in SDDD across simulation runs tended to increase with fragmentation (Fig. 2i).

Finally, increasing habitat amount led to consistently higher CWMDD, species richness, and SDDD (Fig. 2j-l).

### Two-variable interactions

We next examined the effects of simultaneously varying two environmental variables to assess their potentially interacting effects. Here, we first present the findings for simultaneous variation of disturbance and environmental autocorrelation in continuous landscape to investigate environmental variability in the absence of landscape modification (Fig. 3). Second, we assess simultaneous variations of habitat amount and fragmentation in modified landscapes to represent land-use change variables (Fig. 4). Results for all other two-variable combinations are provided in the Supporting figures (Fig. S1-4).

**Figure 3.**
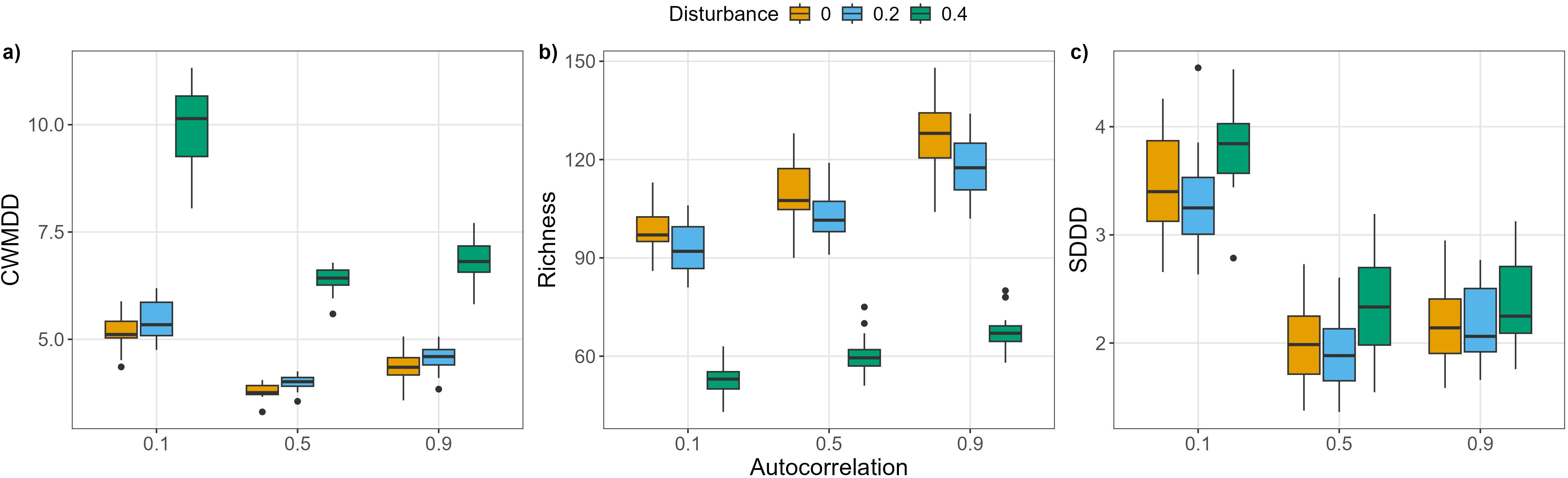
Combined effects of landscape autocorrelation and disturbance, as well as their interactions on three response variables in continuous landscapes – CWMDD (left panel), species richness (middle panel), and SDDD (right panel). Results are based on 20 model repetitions per parameter combination (a total of 180 simulations).

**Figure 4.**
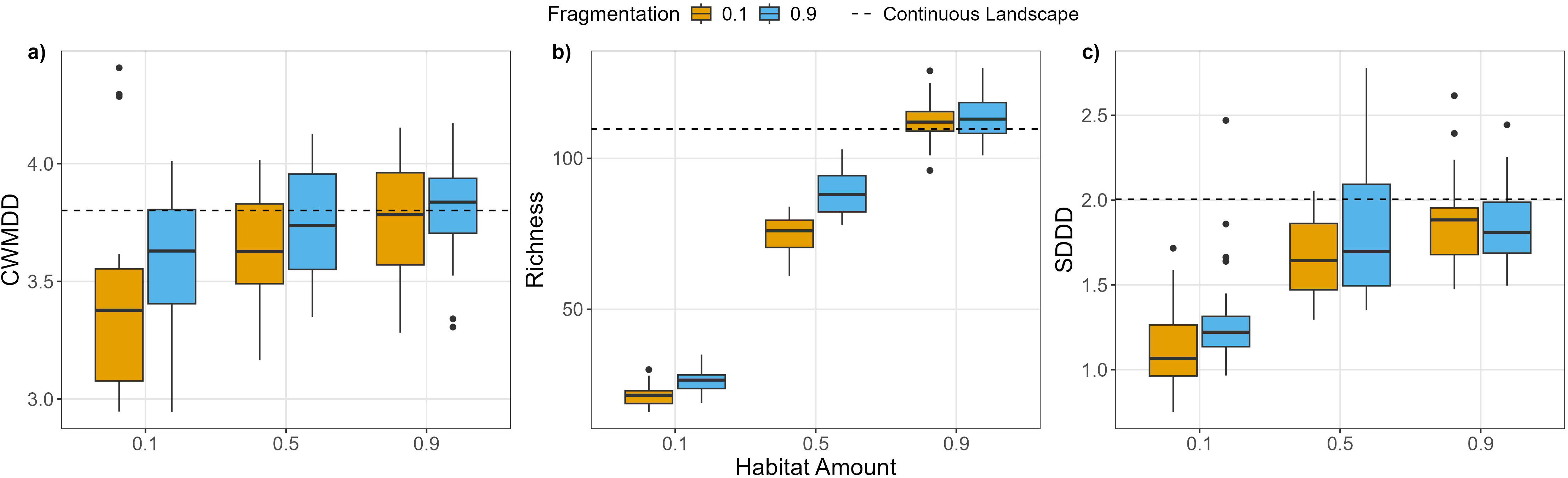
Combined effects of habitat amount and fragmentation level, as well as their interactions, on three response variables in modified landscapes – CWMDD (left panel), species richness (middle panel), and SDDD (right panel). Landscape autocorrelation remained static at 0.5 throughout. Results are based on 20 model repetitions per parameter combination. The dashed line represents the mean results from the continuous experiment (Fig. 3) with no disturbance and an autocorrelation level of 0.5.

When autocorrelation and disturbance were varied simultaneously in continuous landscapes, their main effects did not change compared to the single-variable experiments (Fig. 3). CWMDD increased with lower autocorrelation and higher disturbance, species richness increased with higher autocorrelation and lower disturbance, and SDDD increased with lower autocorrelation and higher disturbance. In addition to the main effects, we found that disturbance increased CWMDD much more at low levels of autocorrelation, while the effect was lower at intermediate and high autocorrelation (Fig. 3a). For species richness, the negative impact of disturbance on richness became slightly stronger at higher levels of autocorrelation (Fig. 3b). SDDD remained consistent with the effects highlighted in the single-variable experiment, and no clear interactions between autocorrelation and disturbance were observed (Fig. 3c).

When habitat amount and fragmentation level were varied simultaneously, the individual effects of the variables again mostly reflected those seen in the single-variable experiments (Fig. 4). All three variables increased with habitat amount, while fragmentation had weaker effects. As opposed to the single-variable experiment, increased fragmentation level did seem to cause a slight increase in CWMDD and the effect was more pronounced under low habitat amounts (Fig. 4a). Also, the positive effect of fragmentation on species richness was stronger at intermediate levels than at low or high levels of habitat amount (Fig. 4b). Across all three metrics, as expected, results under high habitat amounts closely matched those from the continuous landscape experiment (Fig. 3).

### Three-variable interaction

In the following experiment, we explored the effects of varying three environmental variables simultaneously in a modified landscape on the response variables (Fig. 5 – 7). We focused on environmental autocorrelation, disturbance and habitat amount, but neglected fragmentation because of its weak effects in the previous simulation experiments.

**Figure 5.**
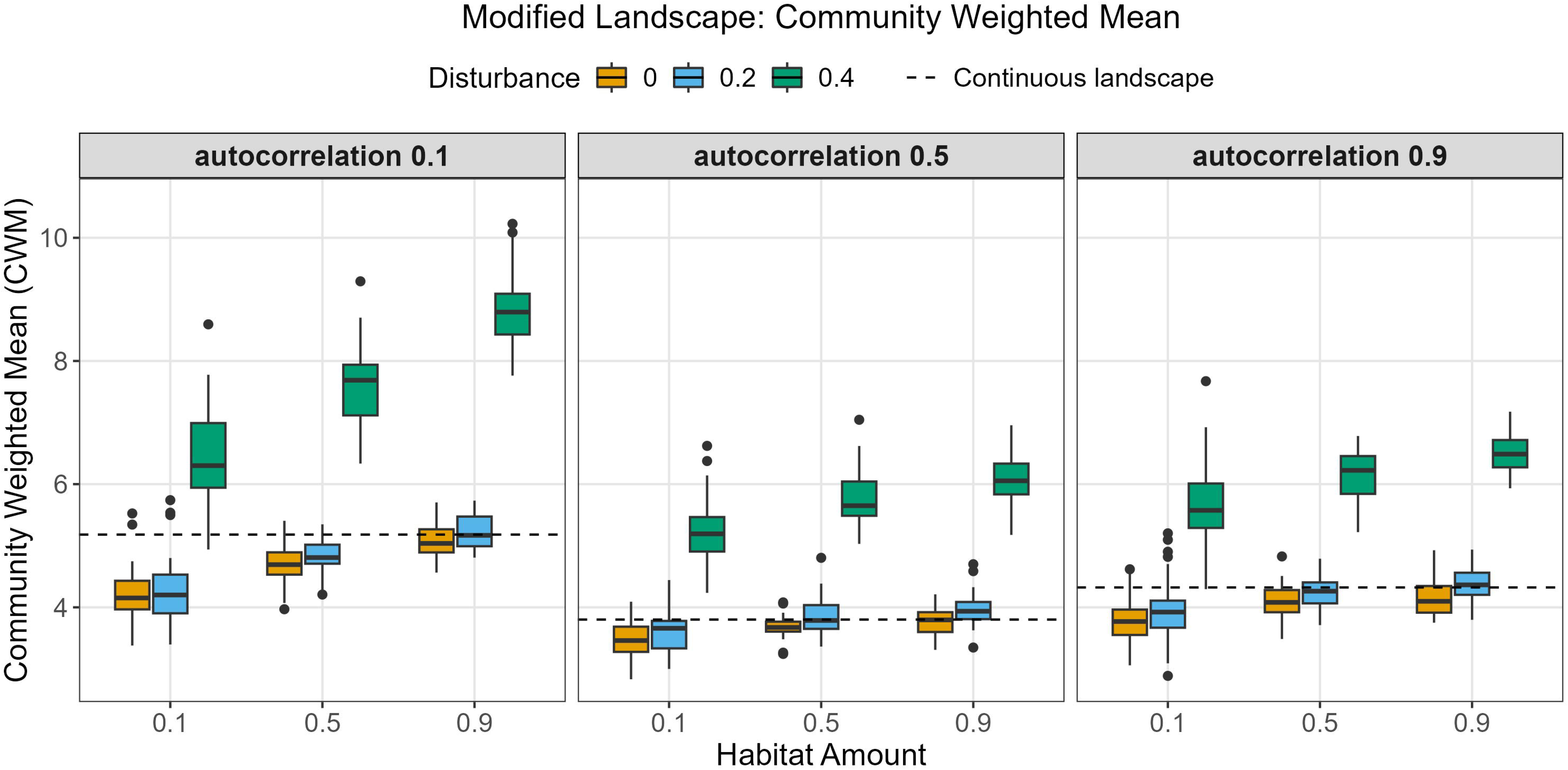
Combined effects of habitat amount, autocorrelation, and disturbance on CWMDD in a modified landscape. Results are based on 20 model repetitions per parameter combination (a total of 1080 simulations). The dashed line represents the average results from the continuous experiment (Fig. 3) with no disturbance at each autocorrelation level.

The effect on CWMDD remained consistent with the single-variable experiments (Figs. 2, 5). The interaction present in Fig. 3, where lower autocorrelation increased the effect of disturbance on CWMDD was also evident here. Additionally, in the modified landscape, we could observe the role that habitat amount plays in this interaction between disturbance and autocorrelation. The interaction was more pronounced in landscapes with high habitat amounts and decreased in landscapes with less habitat. This effect is visible by comparing the differences between the disturbance levels with low autocorrelation across the habitat amount levels. The differences between no and intense disturbance increase with increasing habitat amount, and the effect of disturbance is maximised with high habitat amount and low autocorrelation. Furthermore, in scenarios without or with weak disturbance, the effect of habitat loss on CWMDD was strongly dependent on autocorrelation. With low autocorrelation of the environment, there is a stronger reduction in CWMDD due to low habitat amount compared to the corresponding scenario with high environmental autocorrelation. This happens primarily because with high autocorrelation, CWMDD was already low with 90% or 100% habitat (dashed lines in Fig. 5).

The effects on species richness also remained generally consistent with the single-variable experiment (Fig. 2, 6). The interaction between autocorrelation and disturbance, with increasing disturbance effects at higher autocorrelation, that was observed in Fig. 3, is also present in the current experiment. Additionally, a new interaction emerged between habitat amount and disturbance, where the effects of disturbance on species richness are stronger with increasing habitat amount. We also observed an interaction between habitat amount and autocorrelation, where higher habitat amount in combination with higher autocorrelation increased species richness (Fig. 6).

**Figure 6.**
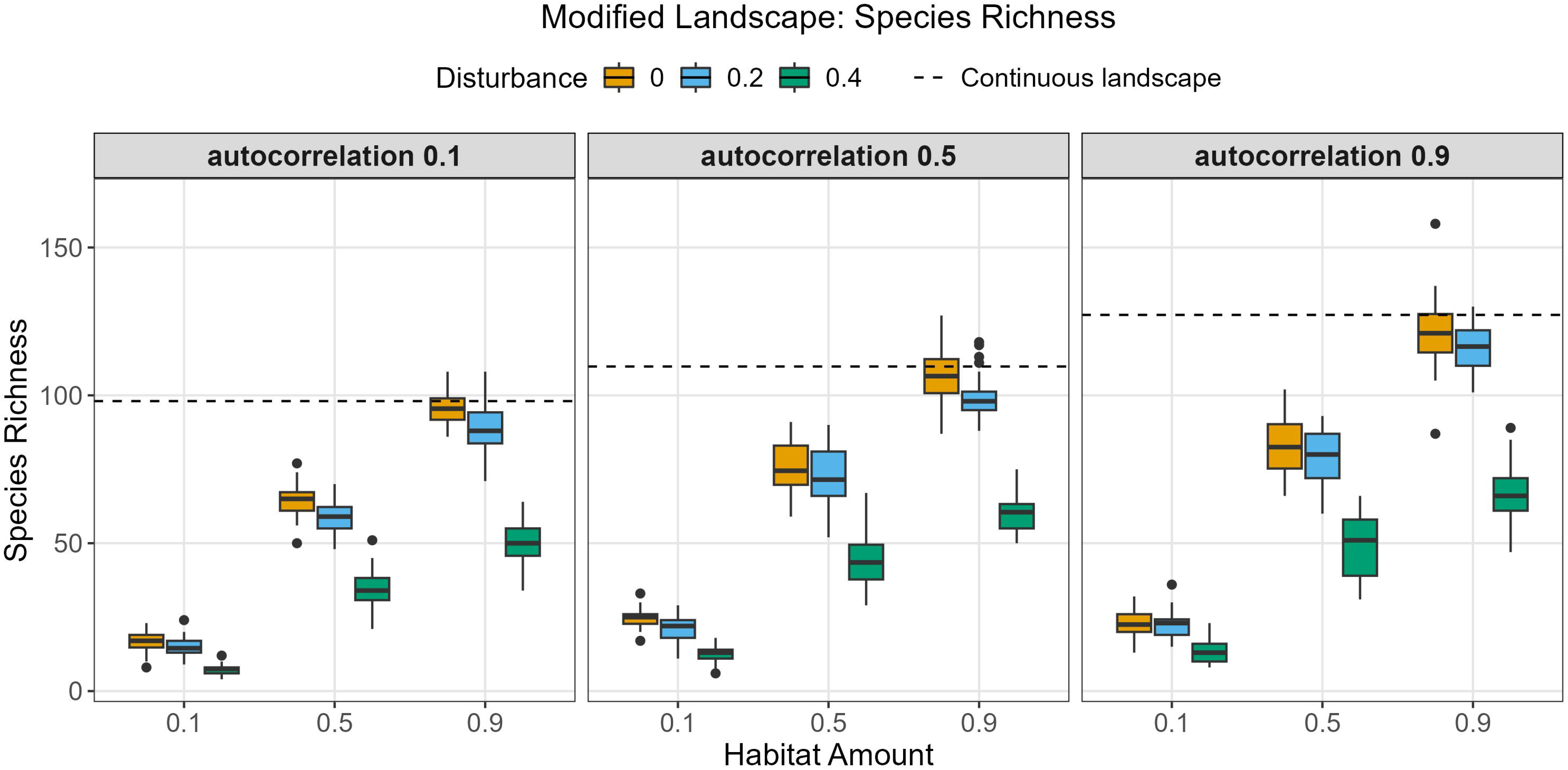
Combined effects of habitat amount, autocorrelation, and disturbance on species richness in a modified landscape. Results are based on 20 model repetitions per parameter combination (a total of 1080 simulations). The dashed line represents the average results from the continuous experiment (Fig. 3) with no disturbance at each autocorrelation level.

Finally, we observed an additional interaction between habitat amount and autocorrelation on SDDD – in low levels of autocorrelation, the positive effects of habitat amount on SDDD were clearly stronger than with intermediate and high autocorrelation (Fig. 7). For all three variables (Fig. 5-7), the effects of modified vs. continuous landscape were consistent with the single variable effects of habitat amount (i.e. continuous landscape results in higher CWMDD, higher species richness, and higher SDDD) as evident by the dashed line representing the continuous landscape.

**Figure 7.**
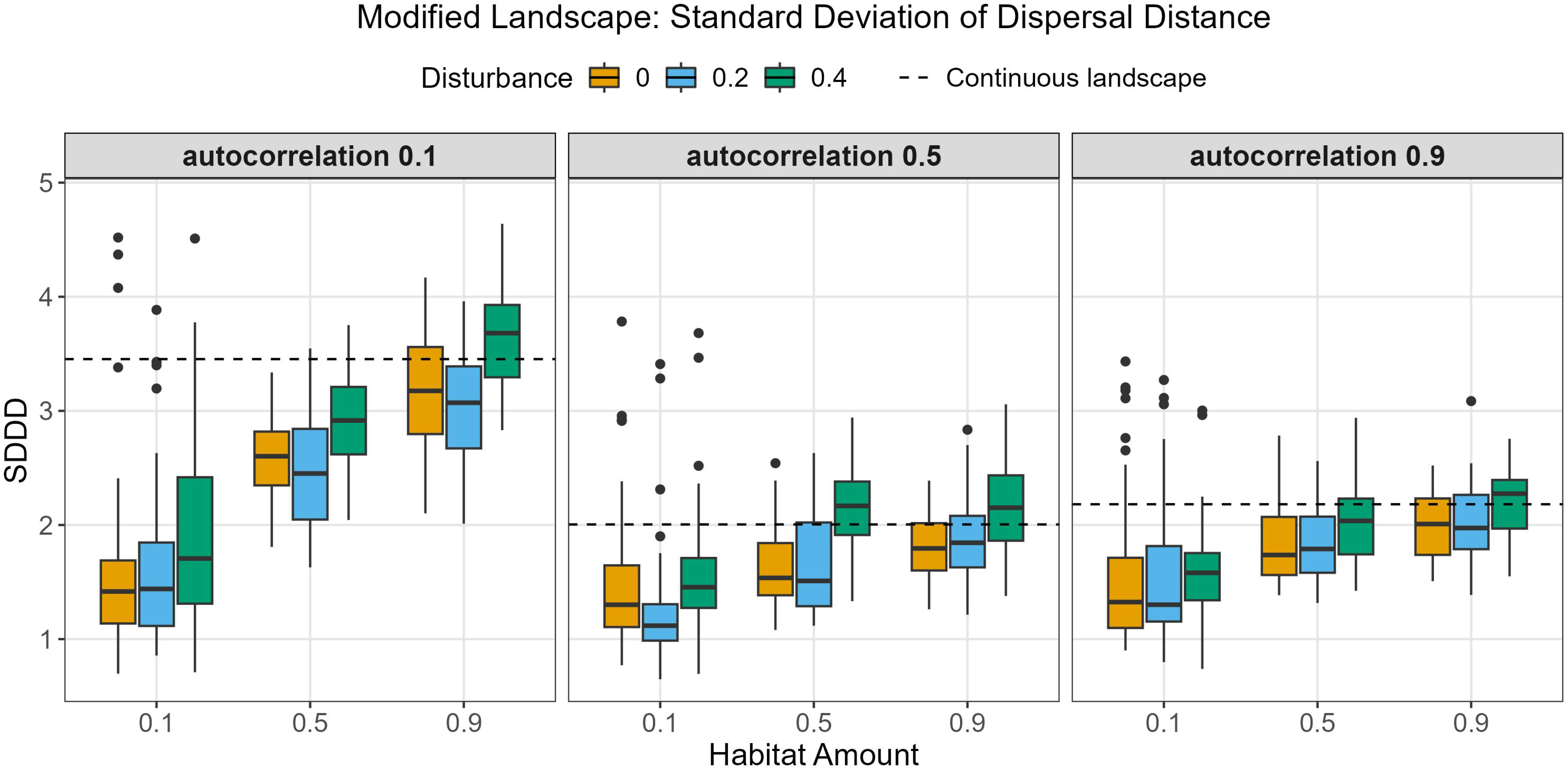
Combined effects of habitat amount, autocorrelation, and disturbance on SDDD in a modified landscape. Results are based on 20 model repetitions per parameter combination (a total of 1080 simulations). The dashed line represents the average results from the continuous experiment (Fig. 3) with no disturbance at each autocorrelation level.

## Discussion

The selection of dispersal strategies in modified landscapes is an essential prerequisite to understanding biodiversity change after land use change. In this study, we aimed at providing new insights into the mechanisms that shape community-level dispersal strategies. Specifically, using an individual-based model, we investigated the independent and interacting effects of four landscape characteristics, namely disturbance, fragmentation, habitat amount, and environmental autocorrelation, on successful dispersal strategies. With this approach, we strive for the synthesis of previous contrasting results, which suggest selection of short (Finand, Loeuille, et al., 2024; Riba et al., 2009; Travis & Dytham, 1999) vs. long dispersal distance (Fronhofer et al., 2014; Thomas, 2000) due to land use change.

We first examined how landscape characteristics shape dispersal strategies in continuous, unmodified landscapes, as understanding selection pressures in these baseline conditions is essential for understanding how landscape modification affects dispersal. In continuous landscapes with a higher spread rate of disturbances, species which exhibited, on average, longer dispersal distances were more successful, as indicated by an increased community-level mean dispersal distance with increasing disturbance spread rate. The observed increase in community dispersal distance under high disturbance likely reflects selection for long-distance dispersal to recolonise disturbed patches, consistent with theoretical predictions and empirical evidence in both plant and animal systems (Clobert et al., 2004; Treep et al., 2021). This is presumably due to the competitive advantage of long dispersers when competing for recolonising disturbed patches. It has to be noted that in our approach, individuals compete for space. Since disturbance creates open space without reducing the habitat quality of this space, species with longer dispersal can access these open resources more easily. Our results also showed that species richness decreased with higher disturbance spread rates, which was expected due to higher mortality and extinction risk.

In continuous landscapes, high environmental autocorrelation selected for short dispersal distances. This is intuitive, as high autocorrelation implies that suitable habitats are more likely to be located near the natal cell. Intermediate levels of autocorrelation selected for even shorter dispersal due to smaller clusters of similar habitats than landscapes with high autocorrelation. Landscapes with a low level of autocorrelation selected for larger dispersal distances, as well as high variation among species, indicating lower selection pressure for species dispersal distances, as indicated by the high variation among species-specific dispersal distances. This can be explained by the high environmental randomness in these landscapes, which do not favour a specific range of dispersal distances. Interestingly, this finding contrasts with previous modelling studies, which reported that highly autocorrelated landscapes favoured greater dispersal distances (Büchi & Vuilleumier, 2012; Travis & Dytham, 1999). This discrepancy can be explained by a key difference in model design: these studies allowed for a philopatric strategy, where individuals could establish in their natal cell. In those models, staying in the natal cell was considered short(er) dispersal, while movement to neighbouring cells was interpreted as long(er) dispersal. In contrast, our model restricted cell capacity to one individual in combination with short propagule longevity, forcing individuals to leave the natal cell to establish elsewhere. The reason for this design choice was to avoid an advantage for short dispersing species by staying in their natal cell (suitable habitat) or dispersing back to it from a nearby similar habitat cell.

Beyond the independent effects of disturbance and environmental autocorrelation on dispersal, we considered the interactive effects of simultaneously varying them in a continuous landscape. We found an antagonistic interaction in the effect of autocorrelation and disturbance on dispersal distances: In landscapes with low environmental autocorrelation, the effects of disturbance on dispersal distances were more pronounced (Fig. 3a). To our knowledge, this antagonistic interaction between disturbance and autocorrelation represents a novel finding. One plausible cause for this effect is the autocorrelated nature of disturbance in our simulations. A disturbed area in a simulation where single species are more aggregated in space due to autocorrelation of habitats will likely be recolonised by the species nearest to the disturbance site. However, in a scenario where individuals are distributed randomly in space (non-autocorrelated landscape), the disturbed patch has a higher chance of being recolonised by long-dispersing species. We also found an antagonistic interaction in the effect of autocorrelation and disturbance on species richness. Higher autocorrelation and lower disturbance led to higher species richness. This could also be explained by the autocorrelated nature of disturbance in our model: Species are more aggregated in space with higher landscape autocorrelation, therefore, local spreading disturbance events are more likely to cause extinction of a specific species.

In modified landscapes, landscape configuration is characterised by the amount and the fragmentation (per se) of the remaining natural habitat. However, we found that the fragmentation level had only weak effects on the selection of dispersal distances. Our results contrast with the results of Hovestadt et al. (2001), who found that increasing fragmentation caused a reduction in dispersal rates. This contrast, again, can likely be explained by the choice to allow philopatric strategy (i.e., recruitment in the site occupied by the mother) in their model, as they showed a strong increase of recruitment within the natal cell with increasing fragmentation. Species richness did increase with increasing fragmentation. This was to be expected as it was previously demonstrated that fragmentation can drive higher species richness under certain conditions, especially due to positive geometric fragmentation effects (Fahrig, 2003; Gelber et al., 2025; May et al., 2019)

The effects of habitat amount on dispersal distances were more pronounced. Lower habitat amount selected for shorter dispersal distances in the community, which is most likely due to an increase in the cost of dispersal – lower amount of nearby habitat cells means higher dispersal mortality due to more dispersal to the matrix. The effect of habitat amount on community dispersal distance was in line with previous studies (Hovestadt et al., 2001). Accordingly, the selection pressure on dispersal distances increased with lower habitat amounts as indicated by reduced mean and variation of species-specific dispersal distances. Species richness increased with increasing habitat amount. The relation between habitat amount and species richness is well established in the literature and corresponds to the well-known species-area relationship (Chase et al., 2020; Fahrig, 2013; Martensen et al., 2012; Rosenzweig, 1995; Watling et al., 2020). The response of dispersal distances to fragmentation and habitat amount is a key finding, revealing that the amount of remaining habitat in a modified landscape is far more influential on community dispersal distances in comparison to the spatial patterns of these habitats. This is in line with general evidence that the amount of habitat has much stronger and more consistent effects on ecological patterns and variables than fragmentation (Fahrig, 2003).

We also explored the interaction between three variables simultaneously – autocorrelation, habitat amount, and disturbance. Beyond the individual effects, our results highlight an interesting three-way interaction between habitat amount, disturbance, and autocorrelation in shaping dispersal strategies. The selection for long-distance dispersal caused by disturbance was most pronounced in landscapes with low autocorrelation and high habitat amount. This three-way interaction can be understood mechanistically: high disturbance creates the need to disperse long distances to escape extinction and/or acquire unoccupied habitat, low autocorrelation requires long-distance travel to find a suitable niche in a random environment, and high habitat amount allows long-distance dispersal by reducing the associated mortality risk. The combination of these three factors creates the strongest selection pressure for long-distance dispersal.

Our finding that less habitat selects for shorter dispersal distances aligns with empirical and theoretical studies predicting reduced dispersal under land-use change (Finand, Monnin, et al., 2024; Riba et al., 2009; Travis & Dytham, 1999). However, other theoretical studies, mainly those grounded in metapopulation theory, have predicted the opposite: that habitat loss should favour longer dispersal (Hanski & Zhang, 1993; Tilman et al., 1994). These contrasting predictions can be reconciled by recognising that metapopulation models typically ignore competition for space, instead focusing on recolonisation of empty patches following local extinctions. They implicitly treat disturbance as the primary mechanism driving local extinction, creating conditions where long-distance dispersal is advantageous for recolonisation. More generally, long-distance dispersal is only beneficial with spatio-temporal variation. That means, when the locations of favourable habitats with available resources and/or low competitive pressure change over both time and space. In classic metapopulation models, stochastic local extinctions, often without reference to a specific mechanism, create this spatio-temporal variability. In contrast, in our model system, spatial variation is caused by environmental variation as well as land use change scenarios, while the additional spatio-temporal variation emerges from spatially-autocorrelated disturbances. When favourable habitats remain static over time, selection favours remaining in suitable habitat patches instead of risking dispersal to hostile locations. Accordingly, our model, which emphasises competition for space in stable habitat configurations at least in the scenarios without disturbance shows different selective pressure than classic single-species metapopulation models.

One interesting implication of our study is that the landscape structure prior to land use change may determine how well the dispersal strategies of communities are adapted to subsequent habitat loss. Specifically, because habitat loss selects for shorter dispersal distances, species from landscapes with high environmental autocorrelation, which also selects for shorter dispersal, may already have dispersal strategies adapted to habitat loss. However, this adaptation may not hold if the modified landscape is also more disturbed than the original continuous landscape. Across our simulations, community dispersal distances were generally lower in modified landscapes compared to continuous ones, as expected given the increased dispersal costs from higher mortality when dispersing to the matrix (Finand, Loeuille, et al., 2024; Travis & Dytham, 1999). Yet when disturbance was introduced to modified landscapes, community dispersal distances became higher than in non-disturbed continuous landscapes, demonstrating that disturbance-driven selection for long dispersal can override habitat loss-driven selection for short dispersal. Under such conditions, the opposing selective pressures (disturbance favouring long dispersal and habitat loss favouring short dispersal) create a novel selective environment to which pre-modification communities are poorly adapted. More broadly, this suggests that the vulnerability of communities to land-use change depends not only on the nature and extent of modification, but also on how the dispersal strategies shaped by pre-modification conditions match or mismatch the selective pressures of the modified landscape.

As mentioned above, some apparent disagreements between our results and previous modelling studies (Büchi & Vuilleumier, 2012; Hovestadt et al., 2001; Travis & Dytham, 1999) can be attributed to differences in whether offspring are allowed to establish in their natal site. This is partly a technical modelling choice, but it also reflects real variation in the life history of species. Mother replacement for example, is possible when mothers are short-lived (e.g., annual plants) and adult mortality occurs before the establishment of juveniles, or when propagules are sufficiently long-lived to persist until the mother dies. In contrast, mother replacement is impossible when propagules are short-lived, and mothers are long-lived. This raises an intriguing follow-up question: Does propagule and adult longevity influence the selection pressure of land use change on dispersal strategies? This issue will not only be interesting for modelling studies, but could also be addressed by empirical comparisons of dispersal-related functional traits in modified landscapes of taxa with different life history strategies.

In this study, we focused on sessile species with passive and undirected dispersal, which does not capture the full range of dispersal strategies found in mobile taxa. Further exploring the development of dispersal strategies with directed and adaptive dispersal would be an interesting next step. Also, dispersal traits were fixed at the species level, precluding adaptive evolution or plastic responses to environmental change, a mechanism known to influence dispersal strategies (Clobert et al., 2009; Travis et al., 2012). Furthermore, dispersal distance was not linked to other species traits, meaning that potential trade-offs were not implemented in the model. In our model, we also chose a rather simplified version of disturbance where we varied disturbance strength only through the probability of spreading, and all individuals in the disturbed area died immediately. An interesting next step would be to explore whether the intensity and frequency of the disturbance have a different effect on community dispersal distances.

## Conclusion

In general, our study shows that dispersal distances in modified landscapes are jointly and interactively influenced by the landscape structure prior to land use change, the intensity of land use change, primarily indicated by habitat loss, as well as disturbances, which may have been part of natural landscape dynamics or may themselves change following land use change. Specifically, our simulations synthesise previous findings by demonstrating that static and thus predictable spatial variation favours shorter dispersal distances, while only spatio-temporal variations of resources and/or competitive pressures can drive the selection of longer dispersal distances. Since the implications of environmental autocorrelation in the unmodified landscape were rarely investigated in this context, we highlight the finding that different levels of environmental autocorrelation may result in different degrees of adaptation of biological communities to land use changes. Understanding dispersal dynamics is crucial because dispersal acts as a link between different spatial scales, mediates species coexistence, and drives community composition under changing environments. By revealing how environmental factors and their interactions influence the selection of dispersal strategies, our findings advance our understanding of how communities respond to global environmental change.

## Acknowledgements

SG acknowledges funding from the German Research Foundation (DFG) under grant number MA 5962/1-1. We thank Selina Baldauf for her careful review of and constructive comments on the simulation model code. We also acknowledge the HPC Service of FUB-IT, Freie Universität Berlin, for providing computing time.

