## Supplementary_Material for "Disturbance and landscape characteristics interactively drive dispersal strategies in continuous and fragmented metacommunities"

### **S1 Two-variable experiment varying habitat amount and disturbance**

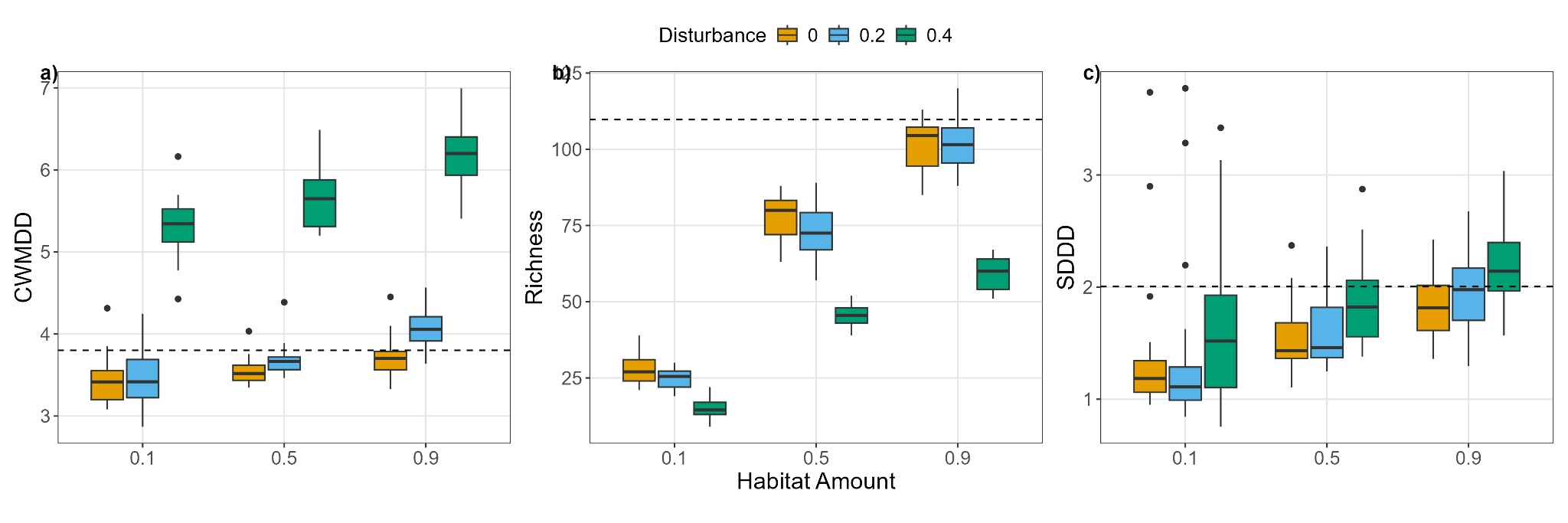

**Figure S1.** Combined effects of habitat amount and disturbance, as well as their interactions, on three response variables in modified landscapes – CWMDD (left panel), species richness (middle panel), and SDDD (right panel). Fragmentation level was held constant at 0.7, and environmental autocorrelation was held constant at 0.5. Results are based on 20 model repetitions per parameter combination.

### **S2 Two-variable experiment varying fragmentation and disturbance**

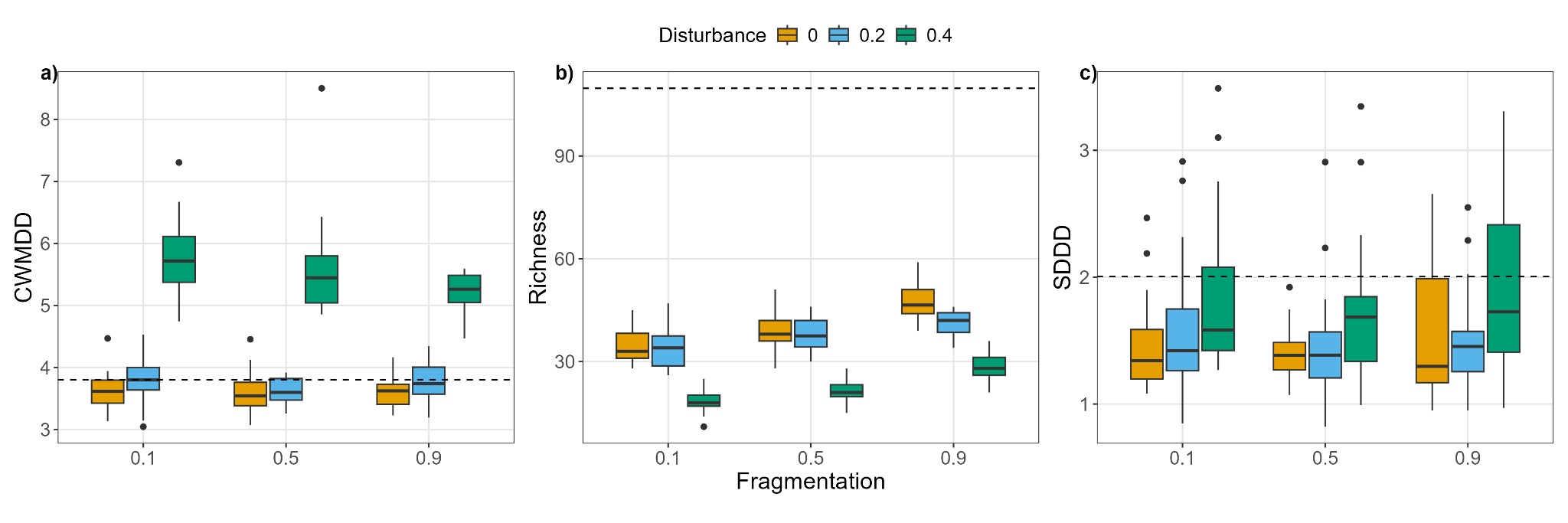

**Figure S2.** Combined effects of fragmentation level and disturbance, as well as their interactions, on three response variables in modified landscapes – CWMDD (left panel), species richness (middle panel), and SDDD (right panel). Habitat amount was held constant at 0.2, and environmental autocorrelation was held constant at 0.5. Results are based on 20 model repetitions per parameter combination.

### **S3 Two-variable experiment varying habitat amount and autocorrelation**

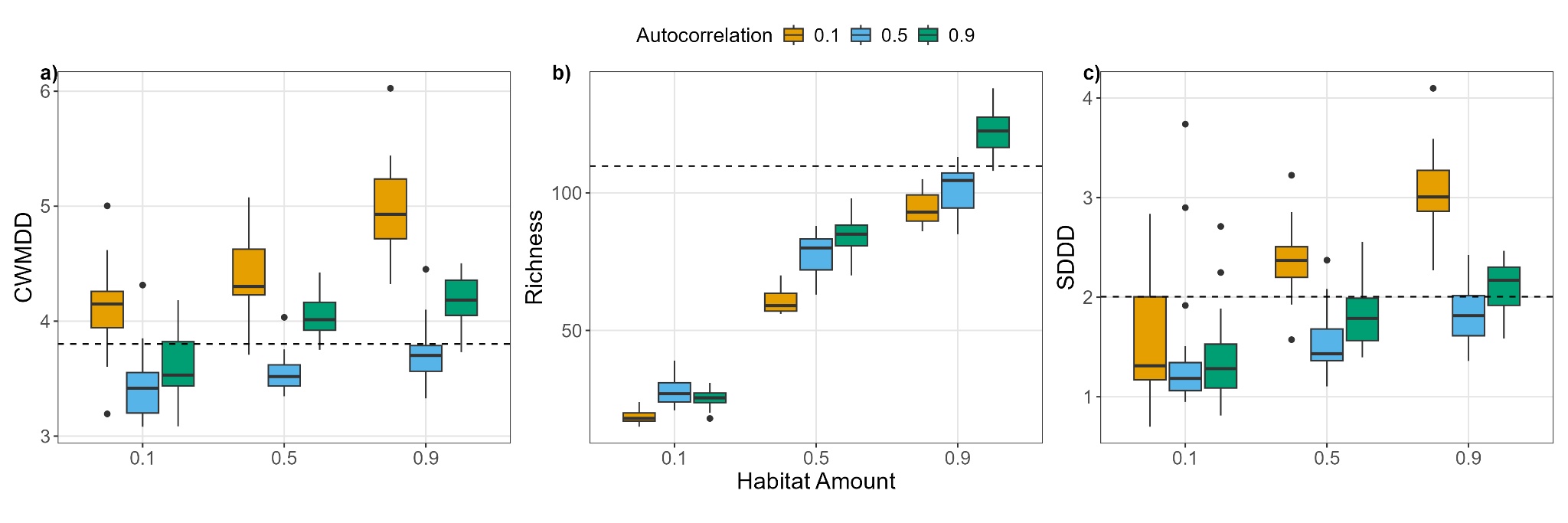

**Figure S3.** Combined effects of habitat amount and environmental autocorrelation, as well as their interactions, on three response variables in modified landscapes – CWMDD (left panel), species richness (middle panel), and SDDD (right panel). Fragmentation level was held constant at 0.7, and no disturbance was applied. Results are based on 20 model repetitions per parameter combination.

### **S4 Two-variable experiment varying fragmentation and autocorrelation**

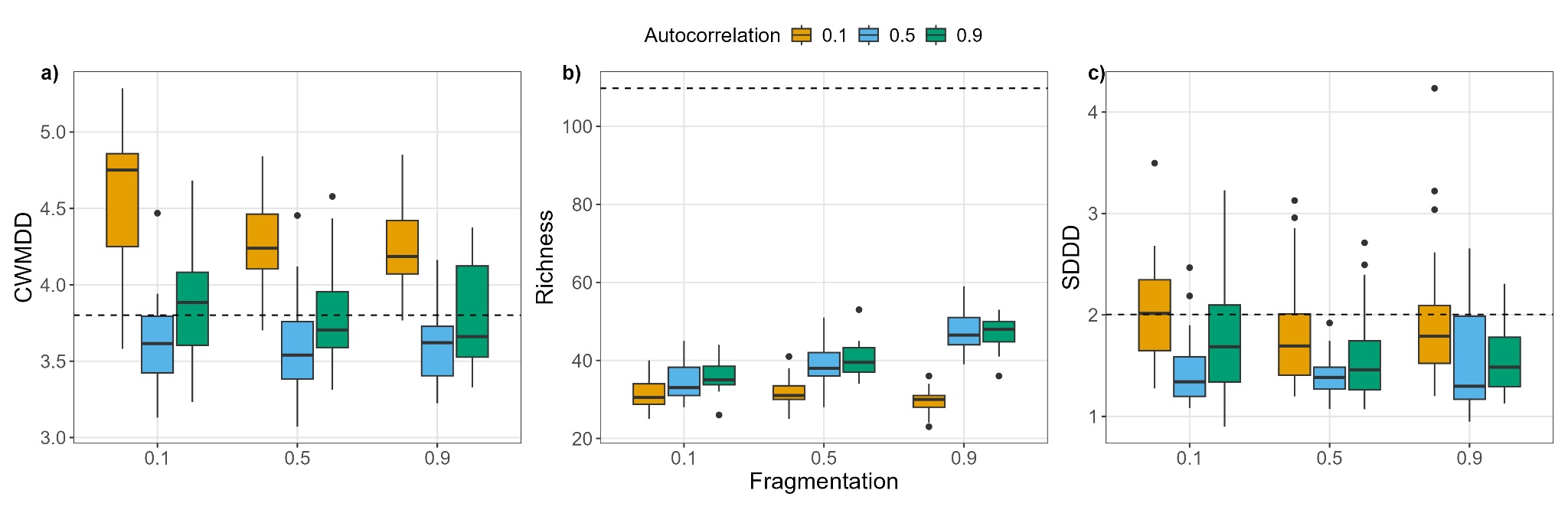

**Figure S4.** Combined effects of fragmentation level and environmental autocorrelation, as well as their interactions, on three response variables in modified landscapes – CWMDD (left panel), species richness (middle panel), and SDDD (right panel). Habitat amount was held constant at 0.2, and no disturbance was applied. Results are based on 20 model repetitions per parameter combination.
